# Enzyme-Responsive Nanoparticles for the Targeted Delivery of an MMP Inhibitor to the Heart post Myocardial Infarction

**DOI:** 10.1101/2022.03.07.483374

**Authors:** Holly L. Sullivan, Yifei Liang, Kendra Worthington, Colin Luo, Nathan C. Gianneschi, Karen L. Christman

**Author notes:** CORRESPONDING AUTHOR FOOTNOTE. These authors contributed equally to this work (Holly L. Sullivan, Yifei Liang).

## Abstract

In this paper, we describe block copolymer amphiphiles consisting of a hydrophilic matrix metalloproteinase (MMP) peptide substrate, and a hydrophobic small molecule MMP inhibitor PD166793 for the treatment of acute myocardial infarction. These resulting drug loaded peptide-polymer amphiphiles (PPAs) assemble in aqueous solution to yield drug loaded micellar nanoparticles. Following minimally invasive intravenous injection, these nanoparticles preferentially exit the vasculature and are physically trapped at the infarcted region of the heart due to MMP-induced peptide cleavage and aggregation. This MMP directed active assembly prevents the material from leaking out into the blood stream, enabling long-term retention. Further, we show that the conjugated MMP inhibitor (PD166793) is inactivated in the core of the micelles and can be released upon the action of proteases and esterases, leading to MMP inhibition. This work establishes a promising targeted nanoparticle platform for delivering small molecule therapeutics to the heart.

## 1. Introduction

Myocardial infarction (MI) is one of the main contributors to the development of cardiovascular disease, the leading cause of global deaths,^1^ and leads to ischemic damage and cardiomyocyte death. During this ischemic injury, the heart enters a pathological state wherein the venous occlusion increases microvascular pressure^2^, resulting in enhanced permeability and retention or “leaky vasculature” in the infarcted region. In addition to this, an upregulation of inflammatory enzymes such an matrix metalloproteinases (MMPs) begin to degrade the native extracellular matrix (ECM) within the heart, compromising mechanical support in addition to cardiomyocyte structure and function.^3^ The enzyme activity is upregulated within hours post-MI and can remain at high levels for weeks to months afterwards.^4^ In the long term, ischemic tissue damage leads to negative left ventricle (LV) remodeling including LV dilation and wall thinning and may eventually lead to heart failure.

Procedures such as coronary artery bypass grafting and catheter-based interventions can revascularize the heart post-MI and many patients will be prescribed anticoagulants, ACE inhibitors, or beta blockers. While these methods can decrease the risk of future adverse cardiac events, they are not addressing the damage that has already occurred. Understanding the limitations of current treatments reveals the clinical need for a therapeutic that can mitigate the extent of injury during the acute phase of MI.

MMP inhibition by systematic administration of pharmacological MMP inhibitors (MMPi) has shown promise in attenuating LV dilatation and reducing infarct size in multiple cardiovascular disease models.^5-7^ However, these drugs face challenges upon clinical translation. First, many of these molecules have poor water solubility and short half-lives (24∼48 hours post administration), and are unable to achieve effective inhibitory concentration in the infarcted heart.^7, 8^ Second, the repetitive dosing and non-targeted systemic delivery has led to off-target side effects, such as joint pain and stiffness associated with musculoskeletal syndrome (MSS).^9, 10^ To address these problems, hydrogels have been used as controlled delivery platforms for MMPi biologics^11, 12^ to improve targeting. The advantages of hydrogels include good targeting and long-term retention (weeks to months) in the infarct. However, the requirement of injection directly into the injured myocardium prevents their use during the acute MI, given the risk of cardiac rupture and arrythmia early post-MI.^13^

In comparison, nanoparticle-based therapeutics can be administrated via minimally invasive intravenous (IV) injection and extravasate into the infarct through the highly permeable vasculature present acutely post-MI.^14, 15^ Active targeting can also be achieved through nanoparticle surface functionalization, such as incorporating substrates for upregulated angiotensin-1 receptor.^16^ Despite this, nanoparticles still face the risk of fast clearance (within hours to days post injection) due to leakage from the infarct and opsonization, wherein biomolecules occlude the surface of particles and subsequently inhibit interaction with tissue receptors.^17^ In light of this, a drug delivery platform that can be administered minimally invasively and then selectively accumulate in the infarct for long-term retention is highly desirable.

Previously, our group developed MMP-responsive nanoparticles that successfully targeted and retained at the infarcted heart for up to 28 days following systematic adiministration.^18^ These materials were made from peptide-polymer amphiphiles where an inert hydrophobic block is followed by a hydrophilic block of MMP-2/MMP-9 cleavable peptides. Following IV injection, these nanoparticles exited the leaky vasculature and were physically trapped in the infarcted region of the heart due to MMP-induced peptide cleavage and aggregation. This MMP directed active assembly prevented the material from leaking out into the blood stream, enabling long-term retention. Building upon this previous work, we conjugated a small molecule MMPi drug, PD166793, to the polymer backbone for enhanced delivery and therapeutic retention in the infarct. PD166793 contains a carboxylate hydrogen that binds tightly to the MMP’s catalytic site^6^, inactivating the enzyme, and has been shown to significantly reduced MMP activity and LV dilation while preserving systolic function in a porcine MI model,^19^ but the requirement of repeated oral dosing to achieve a therapeutic outcome^20^ could increase the risk of negative off-target effects and hinder its clinical translation. We hypothesized that packaging this drug into our enzyme-responsive NPs would allow for targeted delivery to the infarct. When micellar NPs form, the drug will be shielded in the core, then exposed and released following aggregation after enzymatic cleavage of the hydrophilic peptide. As a first step towards the development of this therapeutic, we set out to determine whether conjugation of PD166793 to the MMP responsive nanoparticles would still allow for enzyme mediated aggregation *in vitro* and *in vivo*, and whether the drug would be released and remain active.

## 2. Materials and Methods

### Materials

#### Polymer synthesis and nanoparticle characterization

Organic solvents, including dimethyl formamide (DMF) and diethyl ether were purchased from Fisher Scientific and used without purification. Chemical reagents were acquired from commercial vendors, including Sigma-Aldrich, Thermo Fisher Scientific, and Cambridge Isotope Laboratories Inc, and used as received. The third generation Grubbs catalyst (IMesH_2_)(C_5_H_5_N)_2_(Cl)_2_Ru=CHPh (**G3**),^21^ N-hydroxysuccinimide-functionalized norbornene (NorNHS),^18^ and rhodamine labeled terminating agent (Rho-TA)^21^ were prepared as previously described. Flash column chromatography was performed using silica gel 60 (40-63 μm, 230-400 mesh, 60 A°) purchased from Fisher Scientific. Analytical thin-layer chromatography (TLC) was carried out on silica gel 60G F254 glass plates purchased from EMD Millipore and visualized by observation of fluorescence under ultraviolet light and staining with KMnO_4_ as a developing agent. Dulbecco’s phosphate buffered saline (without Ca^2+^, Mg^2+^) was purchased from Corning. Transmission electron microscopy (TEM) was performed on 400 mesh carbon grids purchased from Ted Pella, Inc.

#### Animal work, histology, and imaging

Female Sprague-Dawley rats were purchased from Envigo. Antibodies were acquired from Dako (anti-α-SMA), ThermoFisher (Alexa Fluor-647), and Vector Laboratories (isolectin). Immunofluorescently stained slides were imaged using an upright Zeiss Fluorescent microscope for 10-20x images and a Zeiss LSM Confocal microscope for 63x magnification images. H&E stained slides were imaged on an Aperio ScanScope CS2 brightfield slide scanner.

#### Cell culture and biocompatibility and MMP activity assessment

Murine L929s were obtained from Millipore. The alamarBlue™ reagent was ordered from Thermo and the MMP-activity assay kit was purchased from Amplite.

### 2.1 Methods

#### Synthesis of PD166793 and functionalized monomer (NorMMPi)

PD166793 was synthesized in four steps from the commercially available 4-bromobiphenyl following modified procedures reported by O’Brien et al. (**Figure 1**, top).^22^ PD166793 functionalized norbornene monomer (NorMMPi) was prepared similarly by coupling L-valine to (N-hexanol)-5-norbornene-exo-2,3-dicarboximide before the establishment of sulfonamide linkage with 4’-bromobiphenyl-4-sulfonyl chloride (**Figure 1**, bottom).

**Figure 1.**
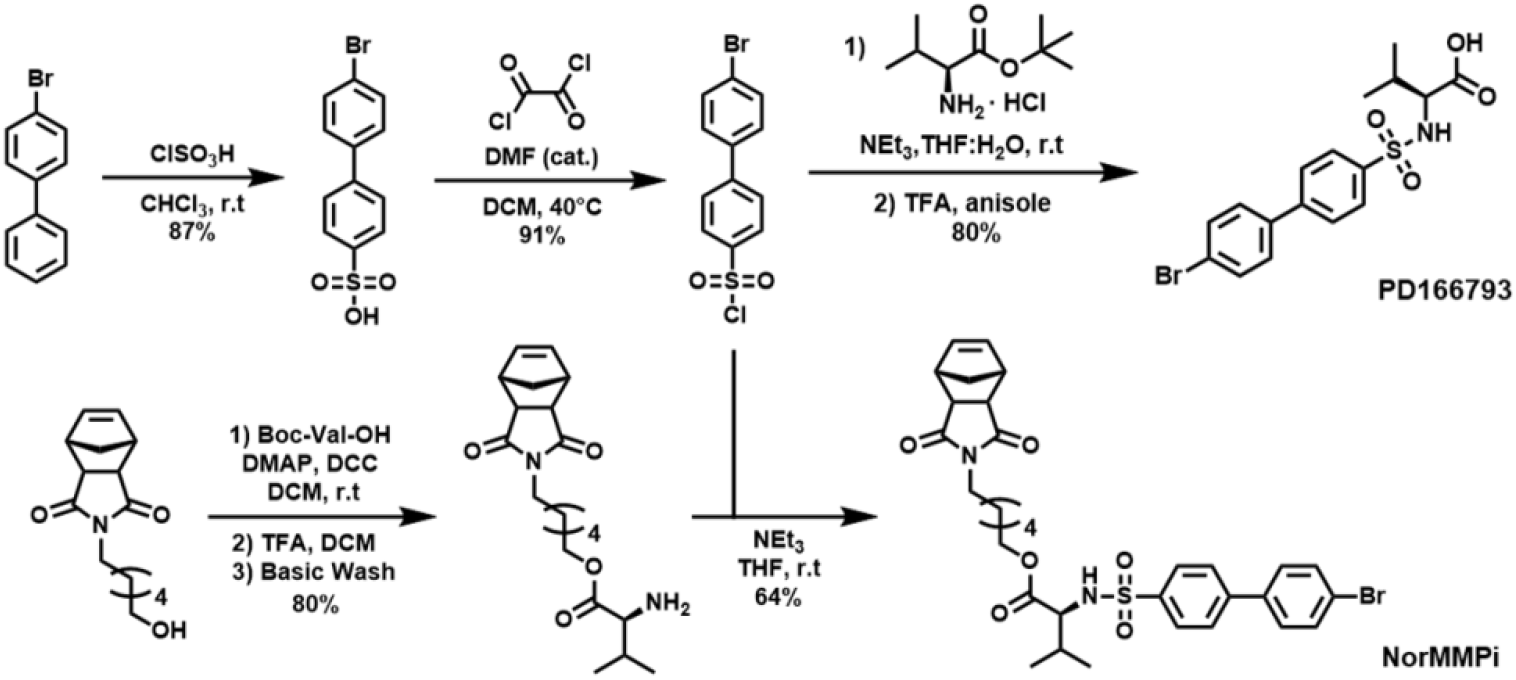
Synthesis of PD166793 and functionalized monomer NorMMPi.

#### Synthesis of MMP-2/MMP-9 cleavable peptide

The MMP cleavable peptide G<u>PLGLAG</u>GWGERDGS (cleavage site underlined) was synthesized via Fmoc-based solid phase peptide synthesis and purified as described by Nguyen et al.^18^ Molar mass of the resulting peptide was confirmed by mass spectroscopy. Detailed procedures can be found in Supporting Information.

#### Synthesis of PD166793 loaded peptide-polymer amphiphile

Peptide-polymer amphiphiles (PPAs) were prepared via graft-to ring-opening metathesis polymerization (ROMP) (**Figure 2A**). To a stirring solution of the third-generation Grubbs catalyst **G3** (IMesH_2_)(C_5_H_5_N)_2_(Cl)_2_Ru=CHPh (1.0 equiv.) in DMF, a mixture of phenyl norbornene (NorPh, 13 equiv.) and NorMMPi (7 equiv.) in DMF was added under N_2_ to yield the random block copolymer NorPh_13_-*co*-NorMMPi_7_. The chain extension was achieved by addition of 5.0 equiv. N-hydroxysuccinimide-functionalized norbornene (NorNHS). The reaction was terminated with a previously reported rhodamine-labeled terminating agent (Rho-TA),^21^ giving rise to the rhodamine-labeled block copolymer. The step growth of polymer molecular weight was confirmed by size-exclusion chromatography coupled with multi-angle light scattering (SEC-MALS) (**Figure S1B, Table S1**). The MMP-cleavable peptide was then conjugated via post-polymerization modification (PPM) in the presence of N,N-diisopropylethylamine (DIPEA). Two equivalents of peptide with respect to NHS was added to promote complete NHS replacement. The peptide consumption was confirmed by the ∼ 50% decrease in free peptide signal on high performance liquid chromatography (HPLC) (**Figure S1C**). The residual peptide was removed through dialysis from DMSO into water and the pure PPAs were collected as the dry powder post lyophilization (**Figure S1B**, *M*_n_ = 13.0 kDa, *Ð* = 1.02).

**Figure 2.**
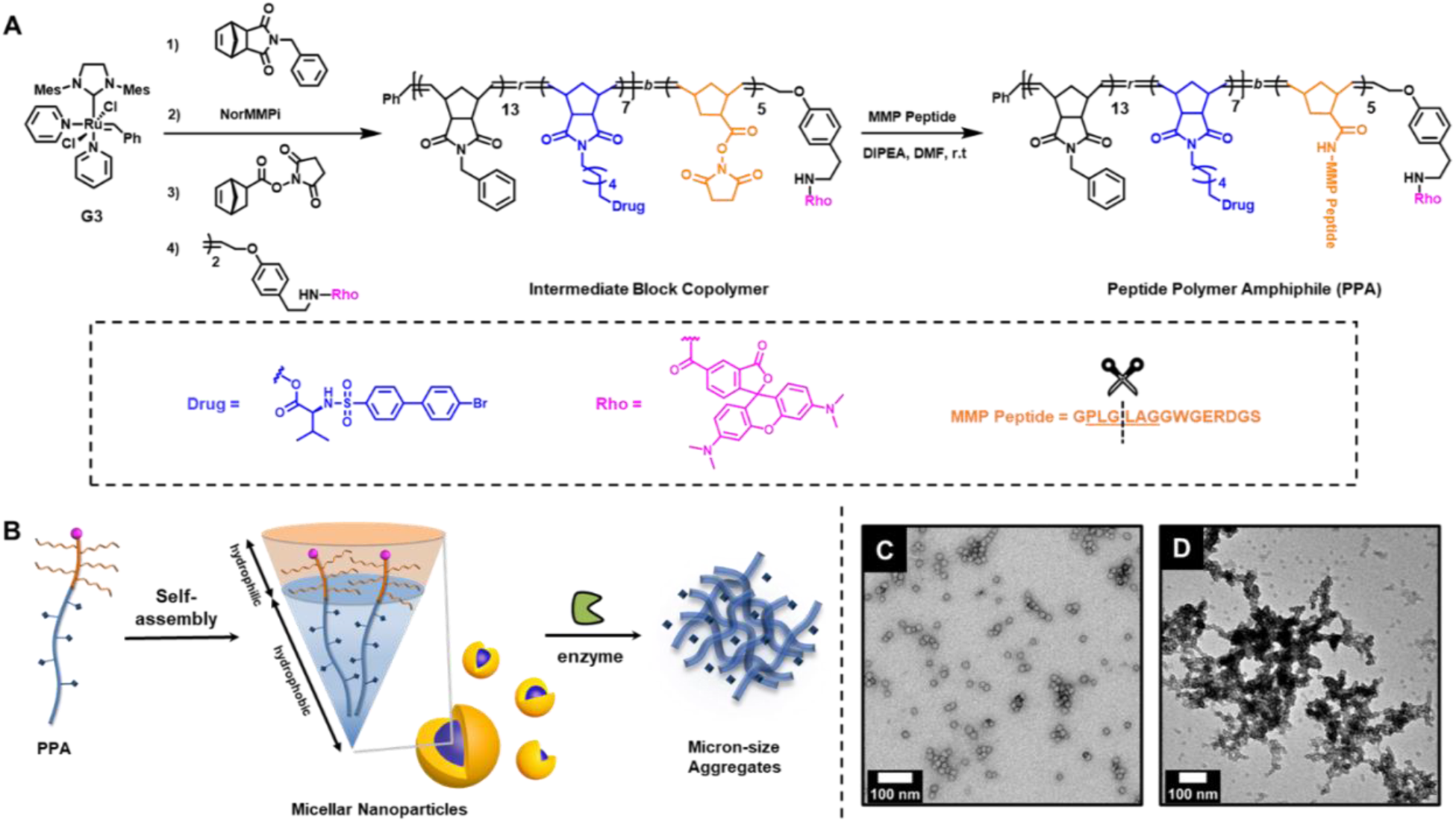
Synthesis of PD166793 loaded nanoparticles and enzyme-induced morphology switch. **(A)** Synthesis of PD166793 loaded peptide-polymer amphiphiles (PPAs). **(B)** Schematic demonstration for PPA self-assembly into nanoparticles and the enzyme-induced microscale aggregates formation. Transmission electron microscopy images of **(C)** nanoparticles and **(D)** aggregates formation post thermolysin treatment.

#### Nanoparticle formulation

PPAs were dissolved at 3 mg/mL in DMSO. 1X Dulbecco’s phosphate-buffered saline (DPBS, without Ca^2+^ and Mg^2+^) was added via a syringe pump at the speed 100 μL/h until reaching 30% DPBS in DMSO (v/v). The solution was left stirring overnight and transferred into SnakeSkin™ Dialysis Tubing (10K MWCO) to dialyze against DPBS for 48 hours with three buffer changes. The resulting solution was filtered through a 0.22 μm PES membrane to remove any large aggregates. The polymer concentration of the filtered solution was confirmed by UV absorbance from rhodamine (**Figure S2**). The nanoparticle solution was concentrated by spin centrifugation to give 300 μM regarding the polymer.

#### In vitro enzyme-induced aggregation

MMPi nanoparticles (NPs) (100 μM, with respect to polymer) were treated with thermolysin, an MMP alternative with improved thermostability, (1 μM) or DPBS for 24 hours at 37 °C in 1X DPBS. The resulting nanoparticle solutions were analyzed by dynamic light scattering (DLS) (**Figure S3**) and transmission electron microscopy (TEM) (**Figure 2C-D**) to examine the change in morphology. For the TEM samples, 5 μL of sample was applied to a 400-mesh carbon grid (Ted Pella, Inc.) that had been glow discharged for 15 seconds. 5 μL of 2 wt.% uranyl acetate solution was then applied and wicked away post 30 sec for staining.

#### *In vitro MMP activity assay* with *NorMMPi*

Due to the poor water solubility of PD166793, stock solution of free drug and NorMMPi (12 mM) were prepared in DMSO. 12 mM ethylenediaminetetraacetic acid (EDTA) in DMSO was also prepared as the negative control due to its strong MMP inhibiting effect by chelating to the Zn cofactor at the MMP active site.^23^ To 1 μL MMP-9 stock solution (38.5 μM stock, 0.5 μM final), 1.5 μL DMSO/NorMMPi/free MMPi/EDTA (12 mM stock, 235 μM final) were added and incubated at 37 °C for 5 minutes. 75 μL MMP peptide (500 μM) in DPBS was then added and incubated at 37 °C for 24 hours before HPLC analysis. The instrument used 0.1% trifluoroacetic acid (TFA) in water as Buffer A and 0.1% TFA in acetonitrile as Buffer B. For each HPLC sample, 50 μL reaction crude was diluted with 150 μL DPBS. 40 μL of the resulting solution was injected and ran over the gradient of 0∼40% Buffer B in 30 min (**Figure S4**). The percent peptide cleavage was quantified and used as the criterion to evaluate MMP activity.

#### Esterase catalyzed PD166793 release

Stock solution of MMPi and NorMMPi were prepared in DMSO at 13 mM. 5 μL esterase (26.4 μM), 5 μL NorMMPi, and 100 μL DPBS were mixed in Eppendorf tubes, giving the final concentration of 1.2 μM esterase: 600 μM NorMMPi. The solution was incubated at either 25 or 37 °C for 24 h. MMPi and NorMMPi at 600 μM in DPBS were also prepared from the stock as the controls. The samples were diluted to 220 μL for HPLC analysis (**Figure S5**). Condition: 10 μL injection, 40∼80% Buffer B over 30 min.

#### Surgical procedures and IV injection

All procedures in this study were approved by the Committee on Animal Research at the University of California, San Diego and the Association for the Assessment and Accreditation of Laboratory Animal Care. Female, Sprague Dawley rats (225 – 250g) underwent ischemia-reperfusion (IR) procedures via left thoracotomy and temporary occlusion of the left anterior descending artery for 35 minutes.^24^ One day post-MI, animals were anesthetized using isoflurane and arbitrarily assigned to IV injection of either 1 mL of MMPi NPs (300µM) or saline and harvested at one day (n = 2) or 6 days (n = 3) post-injection. Animals were euthanized via overdose of pentobarbital (200 mg/kg) and the heart, kidney, spleen, lungs, and liver were collected for histological analysis.

#### Histology and immunohistochemistry

Following euthanasia, hearts were dissected and fresh frozen in OCT for cryosectioning. Hearts were stained with hematoxylin and eosin to visualize the infarcted region of the heart. Slides were scanned on an Aperio ScanScope CS2 brightfield slide scanner. Additional heart sections were stained with anti-α-SMA (1:75 dilution, Dako) and Alexa Fluor-647 (1:500 dilution, ThermoFisher) and isolectin (1:75 dilution, Vector Laboratories) to visualize large arterioles and capillaries.

#### Synthesis of maximally loaded PD166793 nanoparticles and control nanoparticles

Non-fluorescent **PPA**_**Max**_ with increased drug dosage (NorPh:NorMMPi of 0:20) and control **PPA**_**C**_ with no drug incorporation (NorPh: NorMMPi of 20:0) were synthesized similarly as described above (**Figure S6, Table S2**). For both PPAs, the ROMP reactions were terminated with ethyl vinyl ether instead of Rho-TA. They were formulated into nanoparticles (**NP**_**Max**_ and **NP**_**C**_, respectively) via dialysis from DMSO into DPBS as described above. Their morphology before and after incubation with enzyme (i.e., thermolysin) were analyzed via TEM.

#### Cytocompatibility of MMPi NPs and free MMPi

Murine fibroblast cells (L929) were seeded into a 96 well plate and left to adhere overnight. Following cell adhesion, MMPi **NP**_**Max**_ were diluted with sterile PBS to generate concentrations ranging from 41 to 9 µM and were added to the media with PBS and zinc diethyldithiocarbamate (ZDEC) at a concentration of 47mM serving as positive and negative controls, respectively. Treated cells were then incubated for 24 hours before performing an alamarBlue™ assay to evaluate their metabolic activity.

To compare the cytocompatibility of MMPi NPs and free MMPi, L929s were again plated and allowed to adhere overnight. Cells were then treated with MMPi **NP**_**Max**_ or an equivalent concentration of free MMPi ranging from 1200 to 9.375µM. PBS was used as a positive control and DMSO was used to control for the solvent used to dissolve the free drug. After 24 hours of treatment incubation, an alamarBlue™ assay was run on treated cells to evaluate their metabolic activity.

#### Assessment of MMP activity in vitro

Adherent murine fibroblasts (L929s) were plated and allowed to adhere overnight in a 96 well plate at a seeding density of 16,000 cells/well. Media from the plated cells was collected and treated with either MMPi **NP**_**Max**_ (14.28 µM with respect to polymer, 285 µM with respect to drug), free MMPi (285 µM), PBS, or DMSO. The free drug concentration was calculated to match that loaded in the **NP**_**Max**_. Following 20 minutes of treatment, an MMP-cleavage fluorescent FRET peptide (Amplite™) was added to track MMP activity over time following the manufacturer’s instructions. The plate was then incubated at 37 ° C and data points were taken every 20 minutes for 3 hours.

## 3. Results

### Norbornene monomer (NorMMPi) serves as a prodrug and bioactive PD166793 can be released via proteolysis

We first synthesized the MMP inhibitor (MMPi) PD166793 and its functionalized norbornene monomer (NorMMPi). Their inhibitory effects against MMPs were examined using an MMP activity assay. For the control groups, 100% peptide cleavage was observed post DMSO treatment, and no cleavage was detected after EDTA treatment (**Figure S4**). While free PD166793 blocked 70% of MMP activity, no MMP inhibition was observed upon the NorMMPi treatment. This result indicates that NorMMPi serves as a prodrug by shielding the carboxylic acid from interaction with MMP catalytic center, which agrees well with the previous reported drug mechanism of action.^6^ To confirm the bioactive PD166793 can be released from NorMMPi via ester bond cleavage, an esterase mediated proteolysis assay was used. PD166793 has a strong UV absorbance at 270 nm, which enabled us to monitor its release via HPLC (**Figure S5**). While NorMMPi showed good stability in buffer and did not display any HPLC signal under the running condition, we observed complete PD166793 release after 24 hours of incubation with esterase at 37 °C. This result suggests that the drug in the nanoparticle core should be amenable for release upon exposure to the enzyme-rich infarct microenvironment after MMP-induced aggregation.

### PD166793 loaded nanoparticles maintain enzyme responsiveness

PD166793 loaded peptide-polymer amphiphiles (PPAs) (20% wt% drug) were prepared (**Figure 2A**). The degree of polymerization (DP) of each block was optimized, targeting a ratio of 13:7 phenyl-norbornene (NorPh) to NorMMPi to provide a PD166793 plasma level (assuming 100% drug release post 300 nmol PPA injection in the rat model) similar to the effective concentration of 100 µmol/L.^18, 25^ Calculations can be found in Supporting Information.

By solvent switch from DMSO to DPBS buffer, PPAs assembled into spherical micelles with a diameter of 15 nm as visualized by TEM (**Figure 2B-C**). To examine the enzyme responsiveness, the nanoparticles (NPs) were incubated with thermolysin at 37 °C in DPBS buffer (1 µM thermolysin: 100 μM polymer). Both TEM analysis (**Figure 2D**) and DLS measurement (**Figure S3**) confirmed an enzyme-induced morphological switch from nanoparticles to microscale assemblies. Through this *in vitro* analysis, we showed that the PD166793-loaded nanoparticles maintained enzyme responsiveness.

### MMPi NPs localize to the infarct in rat acute MI model

After administration via IV injection, MMPi NPs were visualized in the heart using fluorescence microscopy. As was previously observed with non-drug loaded NPs, MMPi NPs were found to accumulate in the infarcted region of the left ventricle with a majority of the aggregates in the infarct rather than the borderzone (**Figure 3A**).^18^ No MMPi NPs or aggregates were observed in the remote myocardium (septum) or right ventricle. This result is consistent with the non-drug loaded NPs^18^ and demonstrates retained regioselectivity of accumulated NPs in the heart. Like in our previous studies^18^, MMPi NPs were still visible in the heart at 7 days post-injection (**Figure 3B**). At 7 days, we also evaluated the presence of macrophages as an initial evaluation of biocompatibility showing no differences compared to a saline injection (**Figure S7**).

**Figure 3.**
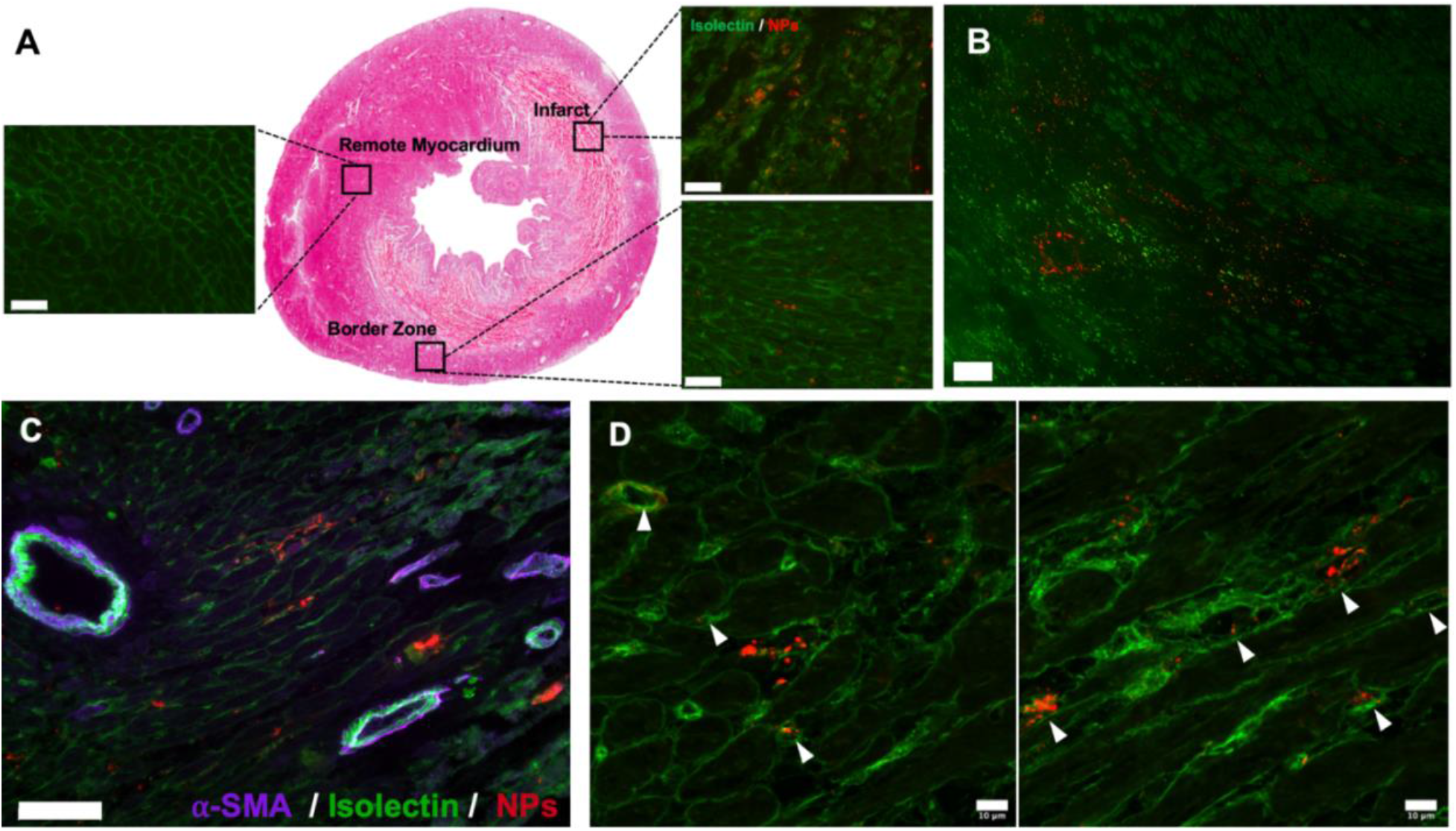
MMPi NPs localize to the infarct and extravasate from leaky vasculature. **(A)** Following IV injection, rhodamine-labeled MMPi NPs localized to the infarct with no accumulation in the remote myocardium. **(B)** At 7 days post-injection, MMPi NPs are still present in the infarct. **(C)** MMPi NPs do not obstruct large vessels (scale bar: 100µm). **(D)** Confocal imaging shows a combination of extracellular deposition of MMPi NPs as well as some aggregate formation within the endothelial cell layer in capillaries (identified by white arrows).

### NP aggregates form in the infarct tissue and not in vasculature

The proposed mechanism of aggregation and accumulation of this platform involves extravasation of nanoparticles from the leaky vessels in the heart post-acute MI. MMPi NPs then will encounter extracellular MMPs and undergo peptide cleavage and accumulation. To confirm localization, smooth muscle cells and endothelial cells were stained with alpha-smooth muscle actin (α-SMA) and isolectin for visualization. MMPi NPs were mostly found outside of the vasculature; fluorescence microscopy images revealed that MMPi NPs do not block arterioles (**Figure 3C**). Through confocal microscopy, we observed MMPi NPs aggregation in the extracellular space as well as in the endothelial layer of some capillaries, which could be attributed to the premature aggregation caused by the MMP released from the endothelial cells^26^ (**Figure 3D**).

### Maximally loading PD166793 does not affect nanoparticle enzyme responsiveness and morphology transition

After confirming that PD166793 loaded **NP** are enzyme responsive and can target infarcted myocardium, we next wanted to examine whether maximally loading PD166793 would affect nanoparticle size and the morphology transition to micron scale aggregates. **PPA**_**Max**_ with a DP of 20:5 (NorMMPi:peptide) was synthesized (**Figure S6A**) as described above and formulated into particles **NP**_**Max**_ (**Figure 4A**). As compared to the original MMPi NPs (20% wt% drug), the drug loading was increased to 40% wt% for **NP**_**Max**_ (detailed calculation in Supporting Information). An inert **PPA**_**C**_ without drug incorporation (NorPh: peptide = 20: 5) was also prepared as a control (**Figure S6B**) and assembled into spherical micelles **NP**_**C**_ (**Figure 4C**). Both NPs underwent aggregation upon thermolysin treatment (**Figure 4B** and **4D**). TEM analysis revealed that both **NP**_**Max**_ and **NP**_**C**_ had similar sizes as the original NPs (15∼20 nm in diameter) and similarly underwent thermolysin induced aggregation to form micro-scale assemblies (**Figure 4B** and **4D**).

**Figure 4.**
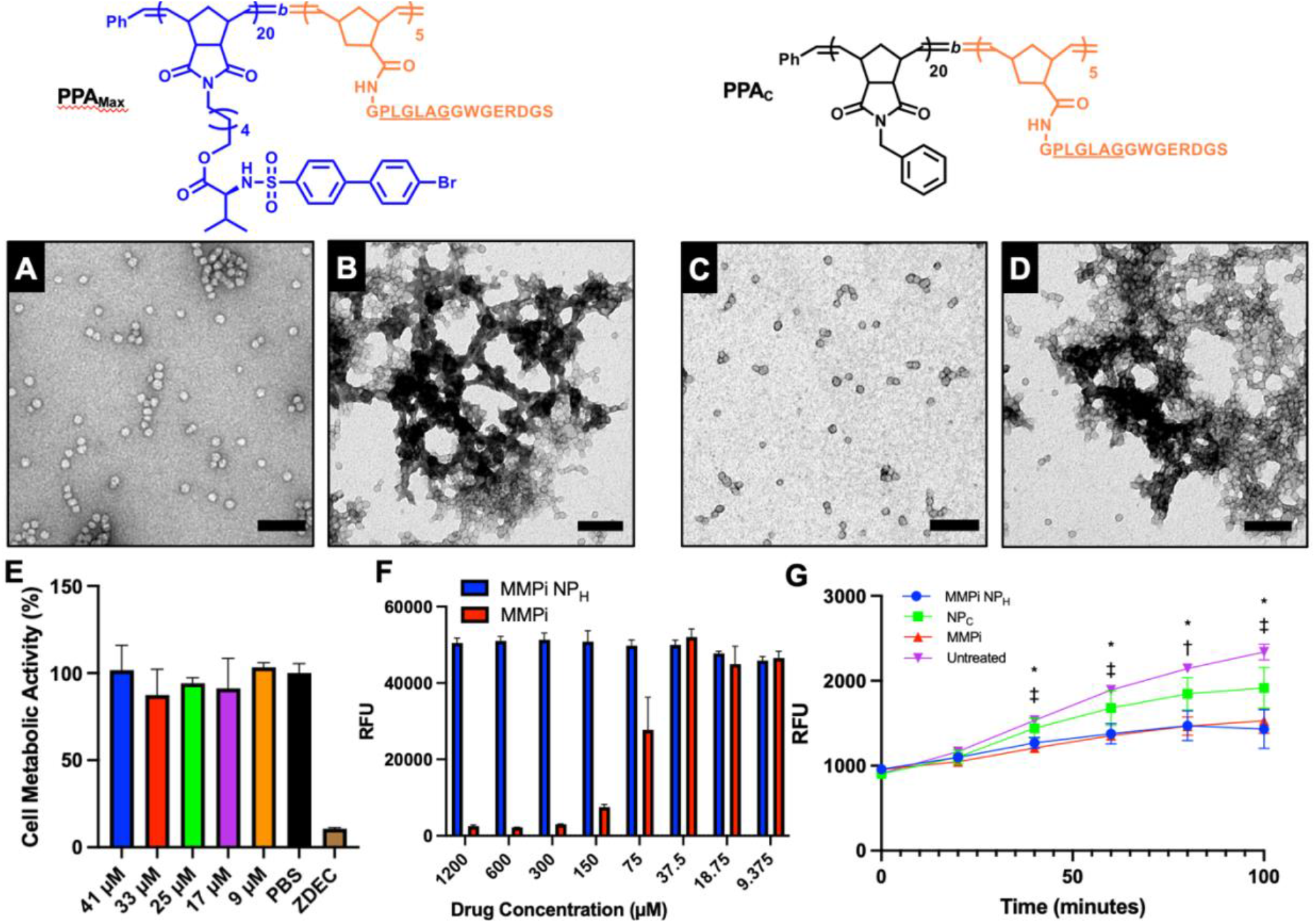
MMPi NP_Max_ and empty NP_C_ are enzymatically responsive and MMPi NPs are cytocompatible and maintain drug bioactivity. Max drug loaded **PPA**_**Max**_ and control **PPA**_**C**_ polymers still form micellar NPs **(A)** and **(C)** of the same size as the initial PD166793 **NP**. Additionally, both still form micron-scale aggregate structures after incubation with thermolysin **(B)** and **(D). (E)** MMPi **NP**_**Max**_ does not significantly impact metabolic activity via alamarBlue™ assay. **(F)** Treatment of L929s with free drug (MMPi) results in reduced metabolic activity at high concentrations whereas conjugated drug (MMPi **NP**_**Max**_) appears to offer a protective effect. **(G)** MMP activity is significantly decreased when treated with MMPi, MMPi **NP**_**Max**_, and even **NP**_**C**_ compared to an untreated control (*P < 0.05 MMPi **NP**_**Max**_ compared to untreated, †P < 0.05 MMPi compared to untreated, ‡P < 0.001 MMPi compared to untreated via 2-way ANOVA with Tukey’s post-hoc test).

### Maximally loaded PD166793 nanoparticles are cytocompatible and bioactive

With the fully loaded MMPi **NP**_**Max**_, we next wanted to evaluate the cytocompatibility of this platform. It has been previously postulated that ∼10% of the injected NPs accumulate in the infarcted region of the heart. In addition, the circulating concentration of MMPi NPs following injection will be ∼18 µM under the assumption that each rat has a total blood volume of 16 mL. This provided us with target concentrations of relevance for our studies. Results from an alamarBlue™ assay show that the concentrations of NPs tested did not significantly alter L929 metabolic activity compared to a positive PBS control up to 41 μM (**Figure 4E**). This result demonstrates preliminary safety of our drug-loaded NP platform. When treated with equivalent concentrations of MMPi **NP**_**Max**_ and free MMPi, we observed a significant improvement in cell metabolic activity for the former at high drug concentrations (75-1200 µM) (**Figure 4F**). This drug concentration correlates to a NP concentration of ranging from 60 to 3.75 µM, the aforementioned physiologically relevant concentration of MMPi NPs in the body and infarcted region. This suggests that a higher dose of PD166793 may be safely tolerated when it is conjugated to the polymeric backbone.

Finally, we sought to confirm the bioactivity of the small molecule MMPi conjugated to the polymer backbone in the fully loaded nanoparticles. In a time-course experiment, we observed that both MMPi **NP**_**Max**_ and free MMPi were capable of significantly inhibiting MMP activity within 40 minutes of incubation compared to a PBS control by 17.4% and 21.3% respectively (**Figure 4G**). From this, we can conclude that packaging the small molecule drug does not negatively impact the bioactivity of PD166793 and that there is rapid drug release from MMPi **NP**_**Max**_.

## 4. Discussion

We have developed a novel enzyme-responsive nanoparticle therapeutic for the potential treatment of acute myocardial infarction (MI). Using ROMP, different dosages of a small molecule MMP inhibitor PD166793 can be incorporated into peptide-polymer amphiphiles though a hydrolysable ester linkage, and subsequently into the core of micellar nanoparticles.

Using a rat MI model, we found that MMPi NPs were able to successfully localize to the infarcted region of the heart following IV injection. Similarly, to the non-drug loaded platform, we observed that MMPi NPs are regiospecific with high accumulation in the infarct, some in the border zone, and little to no accumulation in the remote myocardium. Additionally, these NPs showed prolonged retention in the heart, with aggregates still visible at 7 days post injection. NP retention for this extended window could allow for maximal drug release over time.

In our initial design, we hypothesized that MMP-responsive NPs were extravasating from leaky vasculature in the infarct. Here, we were able to show that arterioles are not occluded by rhodamine-labeled MMPi **NP**_**Max**_. Upon inspection of smaller capillaries using confocal imaging, we observed some nanoparticle presence within the endothelial layer. However, we also see MMPi **NP**_**Max**_ aggregates in the extracellular space. Since it has been previously established that endothelial cells express MMPs,^26^ it is not surprising to observe MMPi **NP**_**Max**_ presence in the capillary endothelium. This could also have therapeutic relevance in inhibiting MMP presence at this location. As further safety evidence, we observed no increase in macrophage presence in the infarct in both saline and MMPi **NP**_**Max**_ -treated animals.

To demonstrate the feasibility of maximally loading PD166793 (**NP**_**Max**_), we first assessed their cytocompatibility *in vitro*. We observed that when the small molecule MMP-inhibitor is conjugated to the polymer backbone, it is tolerated by cells at significantly higher concentrations compared to a comparable concentration of free drug. This is because the drug is not bioactive until it is cleaved from the polymer backbone by endogenous esterase, leading to a slower sustained release of drug compared to a burst release.

When assessing biological activity, we observed equivalent MMP-inhibition by both the MMPi **NP**_**Max**_ and free MMPi *in vitro*. This allows us to conclude that the drug is successfully being released from the polymer backbone and is still bioactive and maintains therapeutic efficacy. Additionally, the comparable onset of inhibition from both MMPi **NP**_**Max**_ and free MMPi implies that the drug release from the polymer backbone is rapid, which could be beneficial for quick therapeutic delivery during the acute phase of MI. Knowing that the drug is still bioactive and that a higher dose may be tolerable when conjugated to the polymer backbone, it is possible that by utilizing our NP platform could allow for safe delivery of high concentrations of PD166793, thus mitigating the need for excessive repeat doses.

This study demonstrates initial proof of concept for the conjugation of a MMPi to MMP responsive nanoparticles. Since any therapeutic that is amenable for chemical conjugation can be included as monomer, we envision this targeted NP platform to be a generalizable approach for drug delivery and the treatment of other inflammatory diseases.

