## Supporting Information for "Enzyme-Responsive Nanoparticles for the Targeted Delivery of an MMP Inhibitor to the Heart post Myocardial Infarction"

AUTHORS EMAIL ADDRESS:

### Experimental Details

#### Synthesis of PD166793

#### 4'-bromobiphenyl-4-sulfonic acid

To a stirred solution of 4-bromobiphenyl (9 g, 38.6 mmol) in chloroform (80 mL), chlorosulfonic acid (5.4 g, 46.33 mmol) was added dropwise at room temperature (r.t). During the addition, precipitation was observed. The reaction mixture was stirred at room temperature for 5 hours. Afterwards, the white precipitate was collected by filtration, washed with cold chloroform and oven-dried at 40 °C to a constant weight to give the final product (10.47 g, 87%). $^{1}$H NMR (500 MHz, Deuterium Oxide) δ 7.84 – 7.76 (m, 2H), 7.61 – 7.54 (m, 2H), 7.51 – 7.45 (m, 2H), 7.40 – 7.33 (m, 2H). $^{13}$C NMR (126 MHz, Deuterium Oxide) δ 142.20, 141.42, 138.04, 131.84, 128.67, 127.07, 126.00, 121.89.

#### 4'-bromobiphenyl-4-sulfonyl chloride

To a suspension of 4'-bromobiphenyl-4-sulfonic acid (5.94 g, 17.5 mmol) in dry DCM (300 mL), oxalyl chloride (11.12 g, 87.6 mmol) and a catalytic amount of dimethylformamide (DMF, 64 mg, 0.88 mmol) was added. The reaction was heated to reflux and stirred overnight. The resulting solution was concentrated to yield a yellow solid, which was dissolved with EtOAc and washed with water. The organic layer was dried with brine and concentrated in vacuo to yield the product (5.2 g, 91%) as a yellow solid. $^{1}$H NMR (500 MHz, Chloroform-*d*) δ 8.13 – 8.08 (m, 2H), 7.80 – 7.75 (m, 2H), 7.68 – 7.61 (m, 2H), 7.52 – 7.45 (m, 2H). $^{13}$C NMR (126 MHz, Chloroform-*d*) δ 147.05, 143.19, 137.42, 132.46, 129.00, 128.02, 127.72, 123.82.

**PD166793 (MMPi)**

To a mixture of L-Valine, t-butyl ester hydrochloride (0.63 g, 3 mmol) and sulfonyl chloride (1.0 g, 3 mmol) in 1:1 THF: $H_2O$ (10 mL), triethylamine ($NEt_3$, 0.61 g, 6 mmol) was added dropwise. The reaction was stirred at room temperature before diluting with EtOAc and washing with 1 M HCl. The organic layer was dried over $MgSO_4$ and concentrated in vacuo. The crude was triturated with hexane and filtered to collect the white solid as the tert-butyl protected PD166793 (1.2 g, 90%). $^{1}$H NMR (500 MHz, Chloroform-*d*) δ 7.92 – 7.87 (m, 2H), 7.67 – 7.62 (m, 2H), 7.62 – 7.57 (m, 2H), 7.45 – 7.40 (m, 2H), 5.13 (d, *J* = 9.9 Hz, 1H), 3.66 (dd, *J* = 9.9, 4.5 Hz, 1H), 2.11 – 1.99 (m, 1H), 1.19 (s, 9H), 1.02 (d, *J* = 6.8 Hz, 3H), 0.86 (d, *J* = 6.9 Hz, 3H).

To a stirred solution of anisole (0.3 g) in trifluoroacetic acid (TFA, 5.6 mL) was added as the tert-butyl protected PD166793. The reaction was stirred for 4 hours at r.t and poured over ice (15 mL). The resulting precipitate was collected by filtration and washed with cold water before dried via lyophilization. The crude was recrystallized via EtOAC/Hex to yield the pure PD166793 as pure white solid. $^{1}$H NMR (500 MHz, Chloroform-*d*) δ 7.91 – 7.84 (m, 2H), 7.68 – 7.63 (m, 2H), 7.63 – 7.59 (m, 2H), 7.49 – 7.43 (m, 2H), 5.11 (d, *J* = 9.9 Hz, 1H), 3.82 (dd, *J* = 9.9, 4.6 Hz, 1H), 2.09 (pd, *J* = 6.9, 4.6 Hz, 1H), 0.96 (d, *J* = 6.8 Hz, 3H), 0.85 (d, *J* = 6.8 Hz, 3H). $^{13}$C NMR (126 MHz, Chloroform-*d*) δ 174.36, 144.56, 138.54, 138.04, 132.26, 128.90, 127.90, 127.44, 123.10, 60.51, 31.42, 19.06, 17.06. ESI-MS: calculated for $C_{17}H_{18}BrNO_4S$ $[M+H]^+$ 412.01 and 414.01; found $[M+H]^+$ 411.9 and 413.9.

**Synthesis of NorMMPi Monomer**

**NorHex-Valine**

To a stirred solution of Boc-Valine (1.5 g, 6.9 mmol) and NorHex (2.0 g, 7.6 mmol) in dry DCM (20 mL) in ice/water bath under $N_2$, a solution of N,N'-Dicyclohexylcarbodiimide (DCC, 1.57 g, 7.6 mmol) and 4-dimethylaminopyridine (DMAP, 169 mg, 1.38 mmol) in 2 mL of DCM was added dropwise. The reaction was allowed to warm to room temperature and stirred for 24 h. After filtration to remove the insoluble urea, the organic layer was washed with 1.0 M HCl, brine and dried over $MgSO_4$. The crude was concentrated in vacuo and purified via column chromatography (40% EtOAc in Hex) to yield a viscous liquid as the Boc protected product (2.8 g, 88%). $^1$H NMR (400 MHz, Chloroform-*d*) δ 6.22 (t, *J* = 1.9 Hz, 2H), 4.96 (d, *J* = 9.1 Hz, 1H), 4.13 (dd, *J* = 9.2, 4.7 Hz, 1H), 4.10 – 3.97 (m, 2H), 3.44 – 3.35 (m, 2H), 3.21 (p, *J* = 1.7 Hz, 2H), 2.61 (d, *J* = 1.3 Hz, 2H), 2.05 (h, *J* = 6.7 Hz, 1H), 1.63 – 1.42 (m, 5H), 1.41 (s, 9H), 1.34 – 1.20 (m, 4H), 1.18 – 1.11 (m, 1H), 0.89 (d, *J* = 6.9 Hz, 3H), 0.82 (d, *J* = 6.9 Hz, 3H).

To 8.5 mL DCM was added NorHex-Boc-Valine (2.8 g, 6 mmol) and TFA (2.8 mL). The reaction mixture was stirred overnight before concentrating in vacuo. The residual was diluted with DCM, washed with $NaHCO_3$, water and brine, and dried over $MgSO_4$ to yield the product (2.0 g, 92%). $^1$H NMR (400 MHz, Chloroform-*d*) δ 6.30 (t, *J* = 1.9 Hz, 2H), 4.11 (td, *J* = 6.6, 2.7 Hz, 2H), 3.53 – 3.40 (m, 2H), 3.35 – 3.21 (m, 3H), 2.69 (d, *J* = 1.4 Hz, 2H), 2.03 (heptd, *J* = 6.9, 4.9 Hz, 1H), 1.70 – 1.47 (m, 8H), 1.45 – 1.28 (m, 4H), 1.23 (dt, *J* = 9.8, 1.6 Hz, 1H), 0.99 (d, *J* = 6.8 Hz, 3H), 0.91 (d, *J* = 6.9 Hz, 3H).

**NorMMPi**

To a mixture of NorHex-Valine **5** (1.2 g, 3.3 mmol) and 4'-bromobiphenyl-4-sulfonyl chloride (1.09 g, 3.3 mmol) in THF (11 mL), triethyl amine ($NEt_3$, 0.67 g, 6.6 mmol) was added dropwise. An immediate formation of white precipitate was observed. The reaction was stirred at room temperature for 5 h before the precipitate was filtered. The filtrate was diluted with DCM and washed with 1.0 M HCl. The organic layer was dried over $MgSO_4$ and concentrated in vacuo to afford white solid as the PD166793 conjugated norbornene NorMMPi (0.9 g, 64%). $^1$H NMR (500 MHz, Chloroform-*d*) δ 7.91 – 7.86 (m, 2H), 7.68 – 7.64 (m, 2H), 7.63 – 7.58 (m, 2H), 7.49 – 7.43 (m, 2H), 6.28 (t, *J* = 1.9 Hz, 2H), 5.26 (d, *J* = 10.0 Hz, 1H), 3.85 – 3.71 (m, 3H), 3.41 – 3.35 (m, 2H), 3.27 (t, *J* = 1.9 Hz, 2H), 2.67 (d, *J* = 1.3 Hz, 2H), 2.05 (pd, *J* = 6.8, 5.0 Hz, 1H), 1.53 – 1.34 (m, 5H), 1.19 (dd, *J* = 7.2, 3.3 Hz, 5H), 0.98 (d, *J* = 6.8 Hz, 3H), 0.88 (d, *J* = 6.8 Hz, 3H). $^{13}$C NMR (126 MHz, Chloroform-*d*) δ 178.11, 178.09, 171.30, 144.30, 138.74, 138.03, 137.83, 132.28, 128.80, 128.00, 127.30, 123.06, 65.39, 61.15, 47.82, 45.18, 42.73, 38.32, 31.70, 28.13, 27.46, 26.32, 25.19, 19.03, 17.41.

**Synthesis of MMP Peptide**

Peptides with the amino acid sequence, GPLGLAGGWGERDGS, were synthesized on rink amide 4-methyl benzylhydrylamine (MBHA) resin via Fmoc-based solid phase peptide synthesis. The underlined amino acids represent an MMP-2 and -9 recognition sequence. The resin was allowed to swell in DMF for 2 hours. Fmoc deprotection was performed by agitating resin in 20% 4-methylpiperidine in DMF for 5 draining, and repeating this procedure for another 15 min. Amino acid couplings were carried out for 45 min per amino acid using N,N,N',N'-tetramethyl-O-(1H-benzotriazol-1-yl)uronium hexafluorophosphate (HBTU) and N,N-diisopropylethylamine (DIPEA) (resin/amino acid/HBTU/DIPEA 1:3:3:6). Final peptides were cleaved from resin by treatment with trifluoracetic acid (TFA), triisopropyl silane (TIPS), dithiothreitol (DTT), and water (TFA/TIPS/DTT/$H_2O$ 88% v/v:2% v/v:5% w/v: 5% w/w) for 2 hours. Peptides were then precipitated in cold diethyl ether and centrifuged at 10000 rpm for 10 min. This procedure was repeated twice. The precipitated peptide was dried in vacuo to give the crude product. The purity of the crude was examined via reverse HPLC running under a mixture of Buffer A (0.1% TFA in $H_2O$) and Buffer B (0.1% TFA in ACN). The peptide afforded a signal at 14 min elution time over a gradient of 15~40% B in 30 min (detection at 214 nm). Same gradient was adopted for peptide purification by a preparation grade HPLC. The purified peptide was dried via lyophilization, giving a white solid. ESI-MS: calculated for $C_{61}H_{94}N_{20}O_{20}$ $[M+H]^+$ 1427.7; found $[M+H]^+$ 1427.7 and $[M+2H]^{2+}$ 714.4. The responsive L-MMP peptide, non-responsive D-MMP peptide, and the cleaved sequence LAGGWGERDGS were both synthesized by this method.

**Representative Procedures for PPA Synthesis**

**Synthesis of Block Copolymer via ROMP**

DMF was freeze-pump-thawed over three cycles. All the monomers and the third-generation Grubbs' catalyst **G3** $(IMesH_2)(C_5H_5N)_2(Cl)_2Ru{=}CHPh$ were weighed separately in vials charged with stirring bars. The monomers and DMF were then loaded into the glovebox under $N_2$. To **G3** (3.28 mg, 1.0 equiv.) in DMF, the mixture of NorPh (14.82 mg, 13 equiv.) and NorMMPi (20.72 mg, 7 equiv.) in DMF was added to afford the first block. After 45 min, an aliquot (~10 μL) was removed for SEC-MALS analysis, and NorNHS (5.3 mg, 5 equiv.) in DMF was added to form the second block. After 1 h, the rhodamine labeled terminating agent (10 mg, 2 equiv.) in DMF was added to quench the catalyst. After 30 min of stirring, the reaction was moved out of the glovebox. The polymers were precipitated in cold diethyl ether and collected by centrifugation at 10000 rpm for 10 min. The same procedures were repeated three times. The final polymers were dried in vacuo and the molecular weights were analyzed by SEC-MALS.

**Peptide Conjugation via Post-Polymerization Modification**

To polymer (46 mg, 1 equiv.) dissolved in DMF, MMP peptide (67.7 mg, 10 equiv. regarding to polymer, 2.0 equiv. regarding NHS) was added, giving a concentration of 50 mg/mL peptide. 5 μL of the solution mixture was removed and dissolved in 245 μL of 15% Buffer B in A for HPLC analysis as the t = 0 reference. DIPEA (30.7 mg, 50 equiv.) was then added to the reaction mixture and stirred at room temperature overnight. After the consumption of peptide was confirmed by HPLC (sample prepared similarly as t = 0), the peptide polymer amphiphiles (PPAs) were precipitated in cold diethyl ether and collected via centrifugation. The resulting crude was dissolved in 1:1 DMSO: water and dialyzed against water in a SnakeSkin™ Dialysis Tubing (10K MWCO). Three water changes were performed in 48 hours and the resulting solution was lyophilized dry to give the pure PPAs. The molecular weight of the dried PPAs was analyzed by SEC-MALS.

Similar procedures were used to prepare the non-fluorescent peptide-polymer amphiphiles with maximum MMPi loading **PPA$_{Max}$** and control **PPA$_{C}$**. Instead of using Rho-TA, the reactions were terminated with ethyl vinyl ether (EVE).

**Calculation for PPA Drug Dosage**

Due to limited water solubility, PD166793 has never been delivered intravenously in an animal model. In a previous report, a plasma concentration of 116 ± 11 μmol/L was shown to attenuate LV dilation and dysfunction in a rat model of progressive heart failure following 4 months of 5 mg·kg$^{-1}$·day$^{-1}$ PO in chow.[1] The following calculation is based on the assumptions:

1) 300 nmol polymer in 1 mL DPBS for injection
2) 250 g average rat body weight with 16 mL of total blood volume
3) 17 mL total body fluid volume (16 mL blood + 1 mL NP solution)

$$\text{Drug wt\%} = \frac{Drug\ weight\ (g/mol \cdot polymer)}{Polymer\ MW\ (g/mol)}$$

$$\text{Drug loading}\left(\frac{\text{mg}}{\text{Kg}}\right) = \frac{Amount\ of\ drug\ (mg)}{Body\ weight\ of\ rat\ (Kg)} = \frac{Amount\ of\ drug\ (mg)}{0.25\ Kg}$$

$$\text{Drug loading}\left(\frac{\mu\text{mole}}{\text{L}}\right) = \frac{Moles\ of\ drug\ (\mu mole)}{Total\ body\ fluid\ volume\ (L)} = \frac{Moles\ of\ drug\ (\mu mole)}{0.017L}$$

**Table 1.** Summary of PPA drug dosage.

| | **Original PPA** | **PPA$_{Max}$** |
|---|---|---|
| **NorPh: NorMMPi (m:n)** | 13:7 | 0:20 |
| **Polymer MW (g/mol)** | 13360 | 20890 |
| **Drug weight (g/mol polymer)** | 2890 | 8240 |

| | | |
|---|---|---|
| **Drug mass (mg/300 nmol polymer)** | 0.87 | 2.5 |
| **Drug moles (μmole/300 nmol polymer)** | 2.1 | 6.0 |
| **Drug wt%** | 20% | 40% |
| **Drug loading (mg/Kg)** | 3.5 | 10 |
| **Drug loading (μmol/L)** | 124 | 353 |

**Macrophage Recruitment Following MMPi NPs Administration**

In animals harvested at 7 days post-MI, heart sections were stained and quantified for general macrophage presence. To identify macrophages, sections were stained with anti-CD68 (1:100 dilution, BioRad) and goat anti-mouse IgG (H+L) conjugate (1:500 dilution, BioRad) and developed using a DAB solution kit (ThermoFischer). Images were taken using a scanning light microscope (Aperio ScanScope CS2). We observed no significant differences in macrophage recruitment between saline and MMPi NPs treated animals. This allows us to conclude that our drug-loaded NPs are well tolerated *in vivo*.

### Supporting Figures

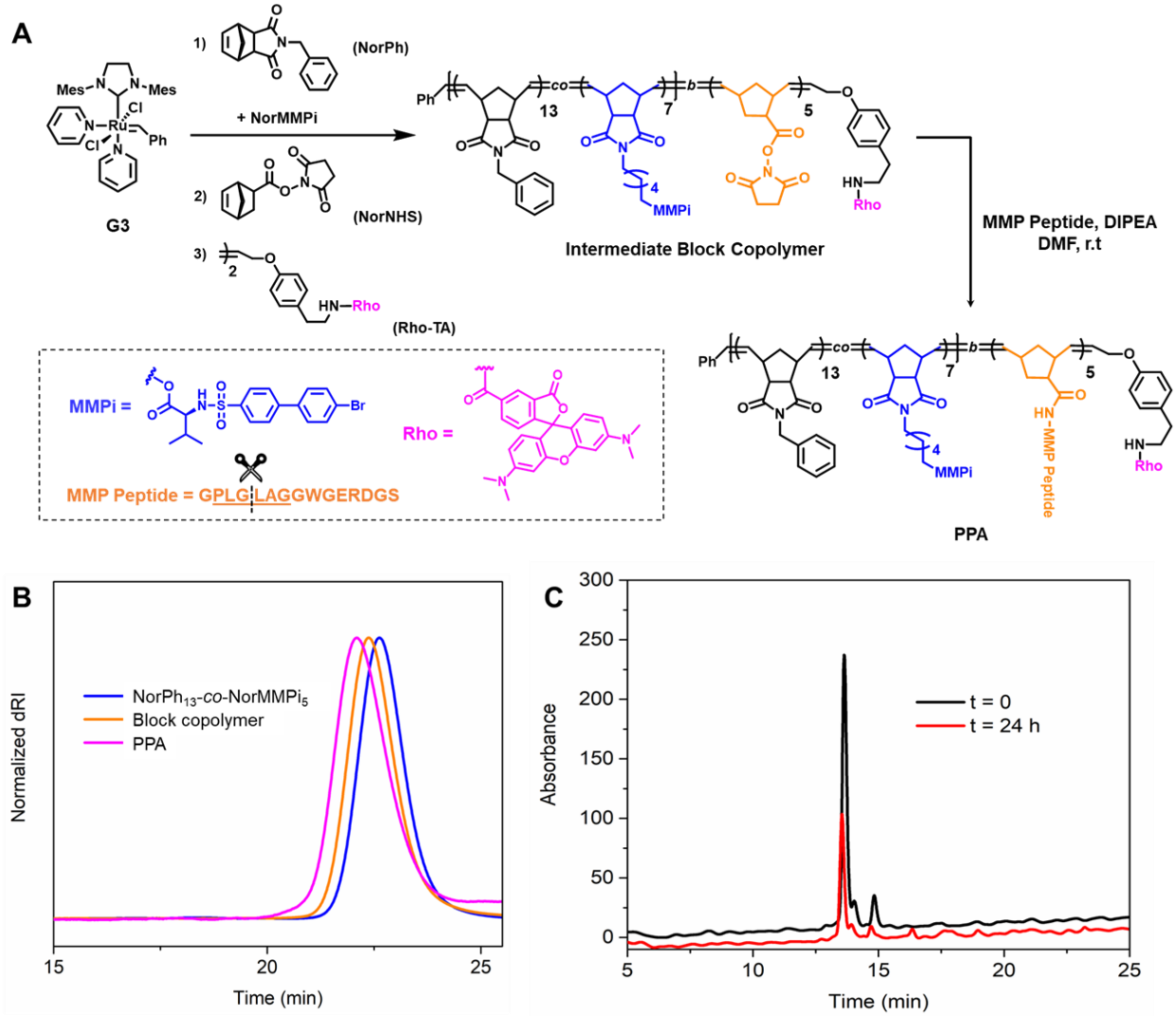


**Figure S1.** Synthesis of PD166793-incorporated peptide-polymer amphiphile (PPA). (A) Reaction scheme for PPA preparation. (B) SEC traces of $NorPh_{13}$-*co*-$NorMMPi_5$, the intermediate block copolymer and PPA post PPM. Polymer molecular weights and dispersities are summarized in **Table S2**. (C) MMP peptide consumption monitored by HPLC.

**Table S2.** SEC-MALS characterization of PPA and intermediates.

| | **NorPh$_{13}$-*co*-NorMMPi$_5$** | **Intermediate block copolymer** | **PPA** |
|---|---|---|---|
| $M_{n,\ theo}$ **(kDa)** [a] | 7.9 | 9.1 | 16.2 |
| $M_{n,\ MALS}$ **(kDa)** [b] | 6.9 | 7.6 | 13.0 |
| ***Đ*** | 1.01 | 1.01 | 1.02 |

[a]Theoretical molecular weight $M_{n,theo} = \sum DP_{monomer} \times MW_{monomer}$.. [b]Molecular weight and dispersity were determined by SEC-MALS with a *dn/dc* of 0.179 mL/g in DMF with 0.05 M LiBr.

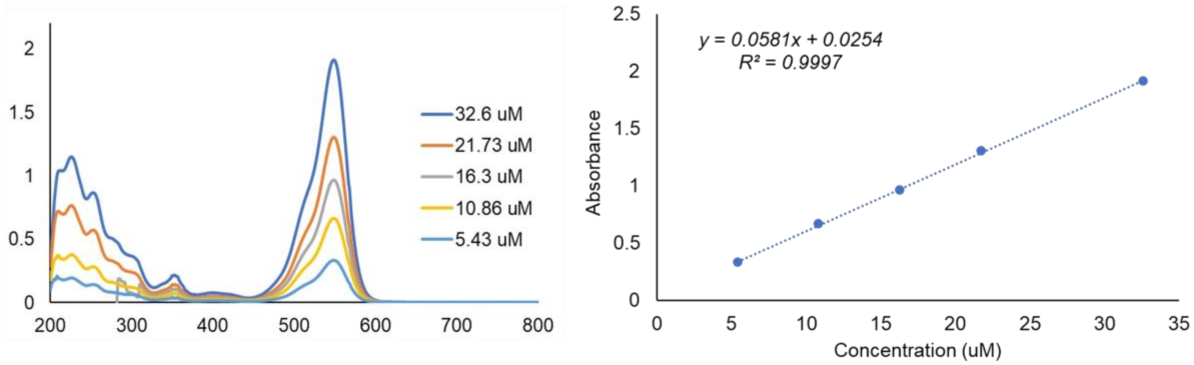


**Figure S2.** UV absorbance of Rho-TA (left) and calibration curve at 548 nm (right).

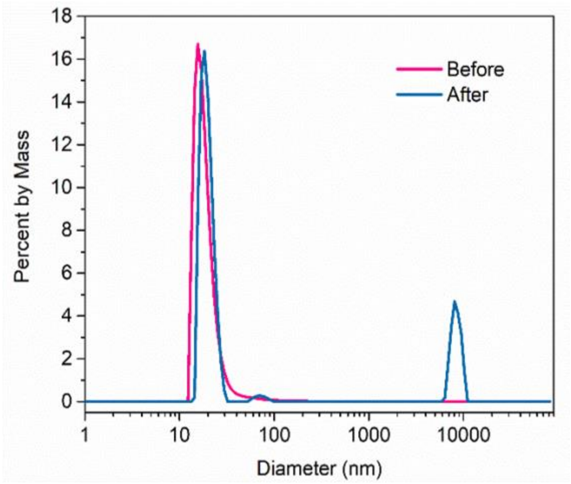


**Figure S3.** DLS analysis of PD166793 loaded nanoparticles (NPs) before and after thermolysin treatment.

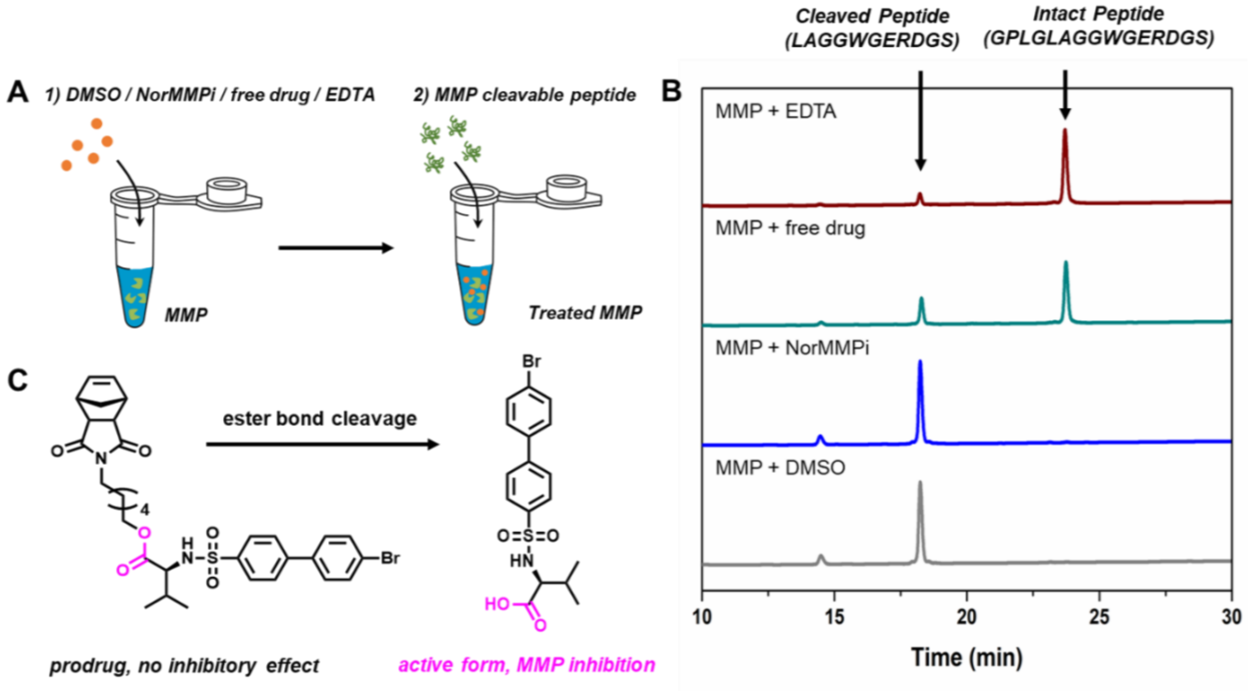


**Figure S4.** MMP-9 activity assay to confirm the drug mechanism of action. (A) Schematic illustration of the experimental setup. (B) HPLC traces that showed peptide cleavage post incubation with MMP-9 treated with DMSO, NorMMPi, free PD166793 or EDTA. (C) PD166793 release from NorMMPi monomer through ester bond cleavage.

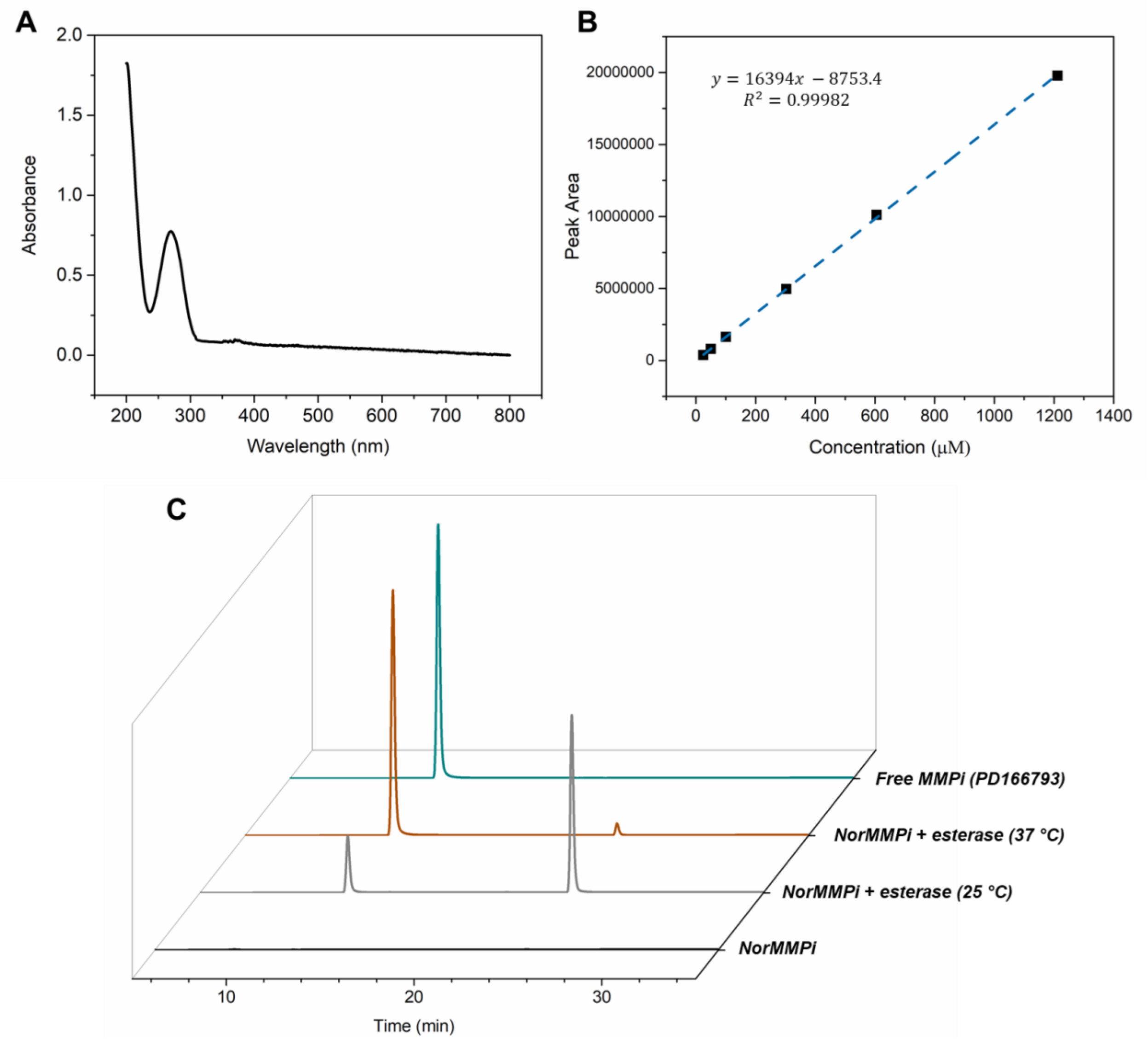


**Figure S5.** Esterase-catalyzed PD166793 release from NorMMPi. (A) UV-Vis spectrum of PD166793 in acetonitrile. Maximum absorbance was at 270 nm. (B) HPLC calibration curve of PD166793. (C) HPLC traces of NorMMPi alone, NorMMPi post esterase treatment at 25 and 37 °C, and PD166793 control.

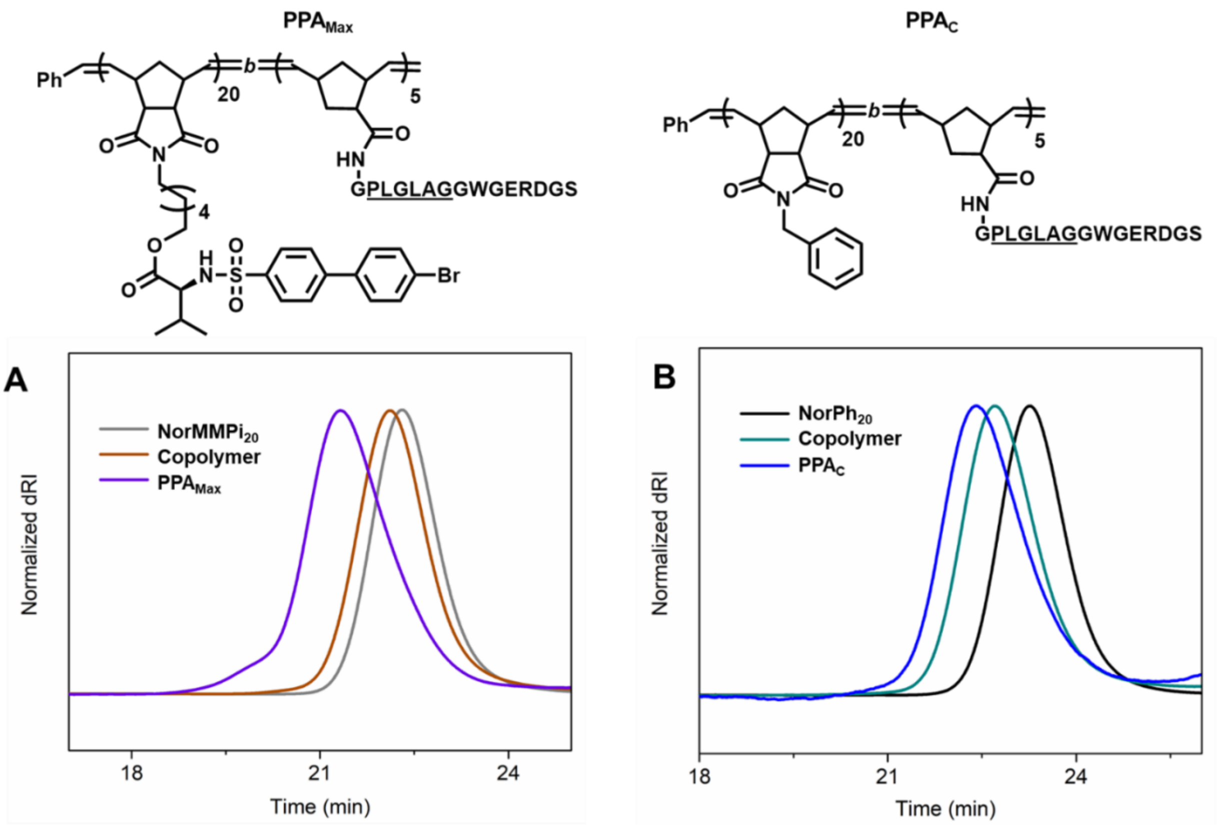


**Figure S6.** Synthesis of **PPA$_{Max}$** and **PPA$_{C}$**. (A) SEC traces of NorMMPi$_{20}$, the intermediate copolymer (NorMMPi$_{20}$-*b*-NorNHS$_5$) and **PPA$_{Max}$.** (B) SEC traces of NorPh$_{20}$, the intermediate copolymer (NorPh$_{20}$-*b*-NorNHS$_5$) and **PPA$_{C}$**. Polymer molecular weights and dispersities are summarized in **Table S2**.

**Table S3.** SEC-MALS characterization of **$PPA_{Max}$**, **$PPA_C$** and intermediates.

| | **$NorMMPi_{20}$** | **$NorMMPi_{20}$-*b*-$NorNHS_5$** | **$PPA_{Max}$** | **$NorPh_{20}$** | **$NorPh_{20}$-*b*-$NorNHS_5$** | **$PPA_C$** |
|---|---|---|---|---|---|---|
| **$M_{n,\ theo}$ (kDa)** [a] | 13.2 | 14.3 | 20.9 | 5.1 | 6.2 | 13.4 |
| **$M_{n,\ MALS}$ (kDa)** [b] | 9.2 | 9.7 | 16.0 | 5.5 | 5.9 | 13.5 |
| ***Đ*** | 1.01 | 1.01 | 1.06 | 1.02 | 1.01 | 1.15 |

[a]Theoretical molecular weight $M_{n,theo} = \sum DP_{monomer} \times MW_{monomer}$.. [b]Molecular weight and dispersity were determined by SEC-MALS with a *dn/dc* of 0.179 mL/g in DMF with 0.05 M LiBr.

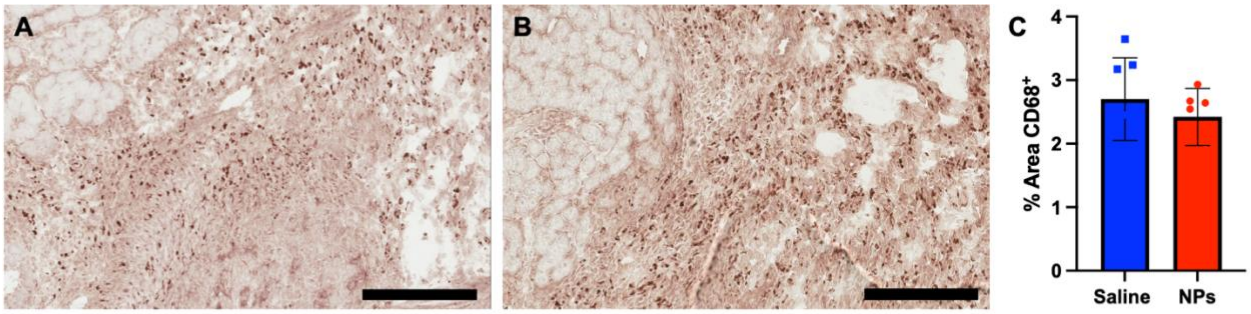


**Figure S7.** CD68 staining in the infarct at 7 days post injection show no sign of increased macrophage presence in the saline-injected (A) or NP-injected (B) hearts (Scale bars are 200 µm). (C) Quantification of these images confirms this trend.

**NMR Spectra**

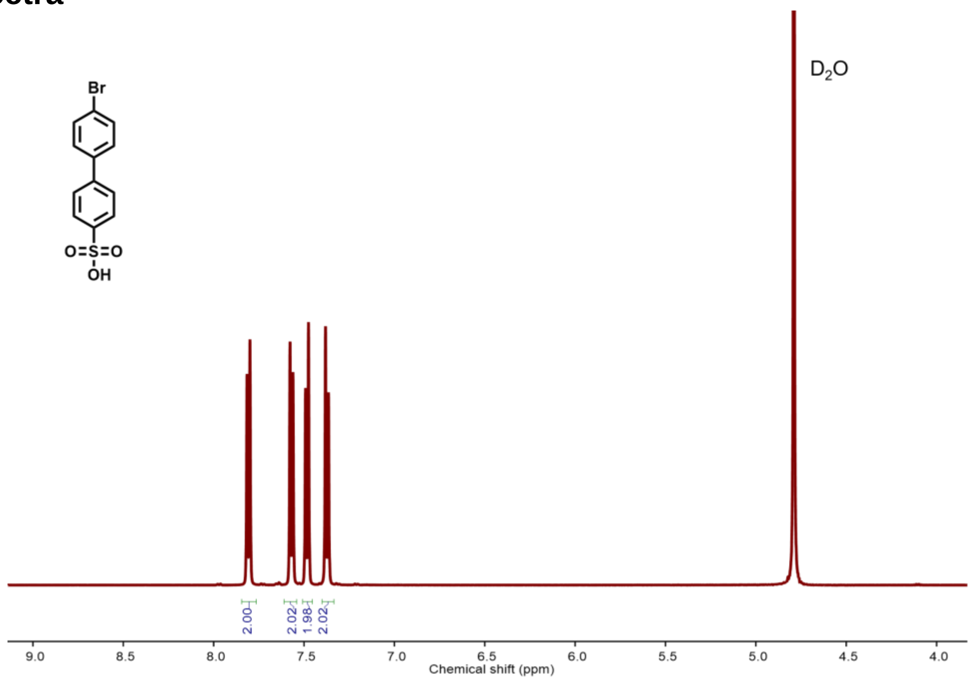


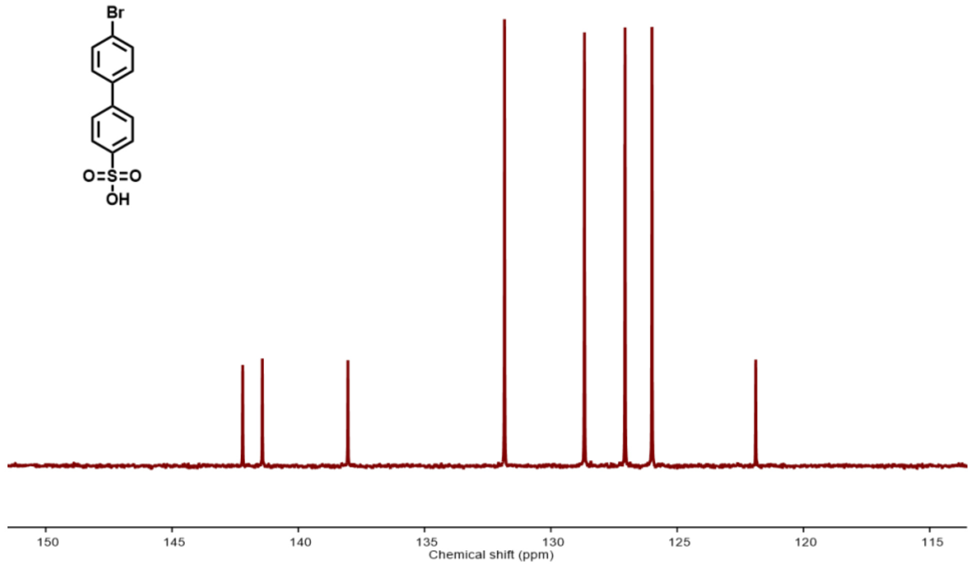


**Figure S8.** $^{1}H$ (top) and $^{13}C$ NMR (bottom) spectra of 4'-bromobiphenyl-4-sulfonic acid in $D_2O$.

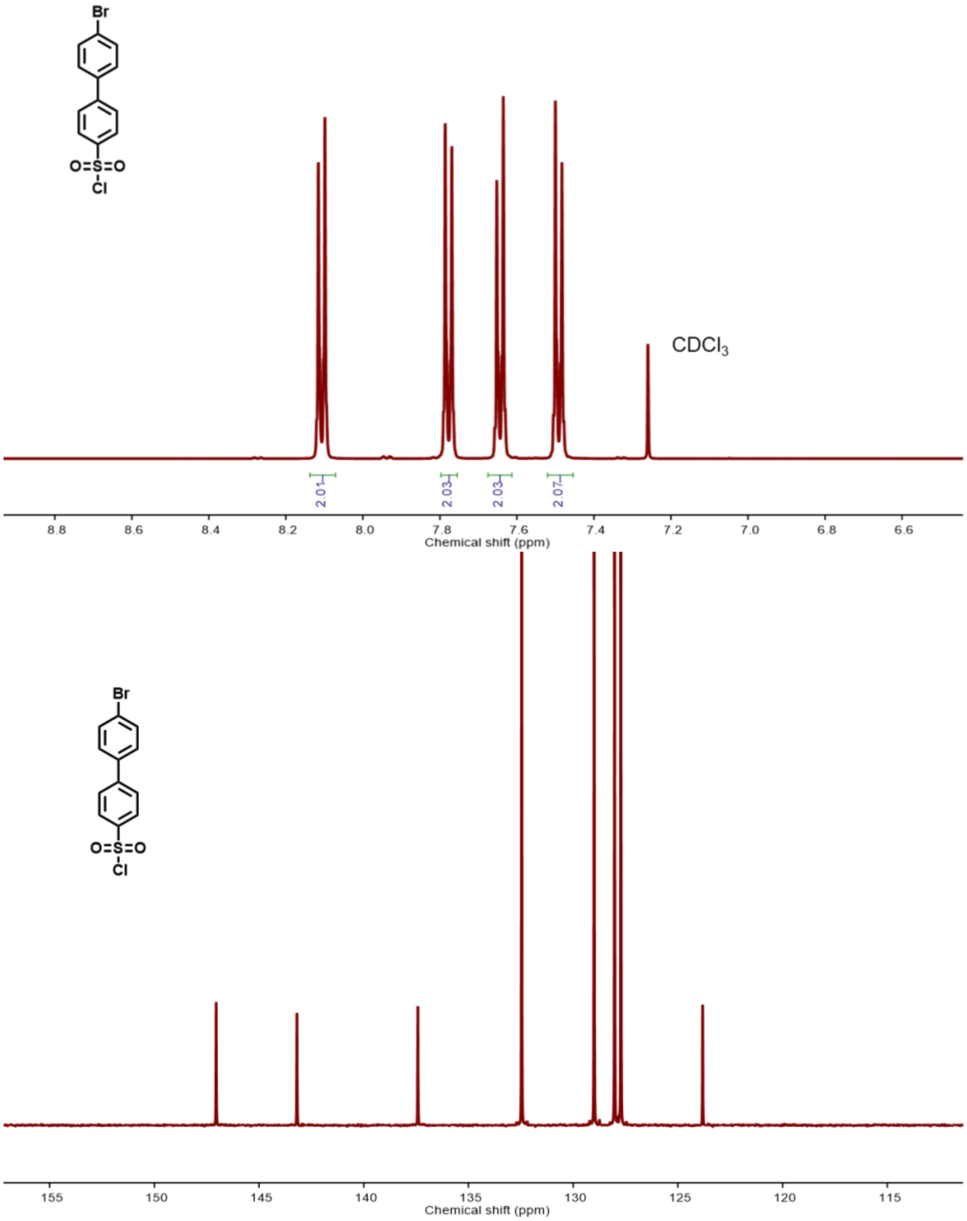


**Figure S9.** $^{1}$H (top) and $^{13}$C NMR (bottom) spectra of 4'-bromobiphenyl-4-sulfonyl chloride in $CDCl_3$.

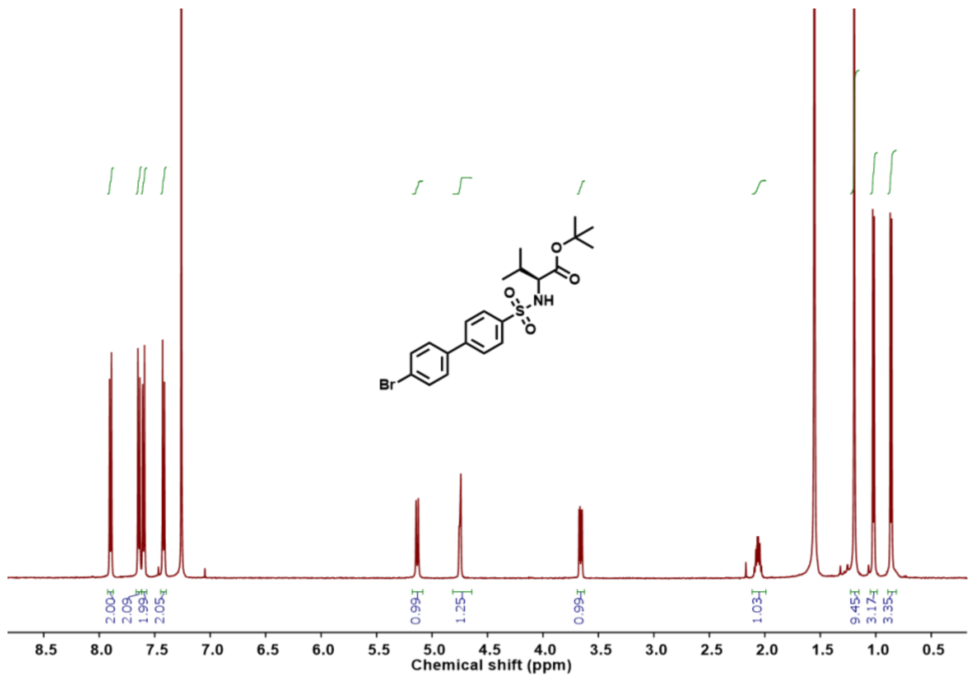


**Figure S10.** $^1$H NMR spectrum of t-butyl ester protected PD166793 in $CDCl_3$.

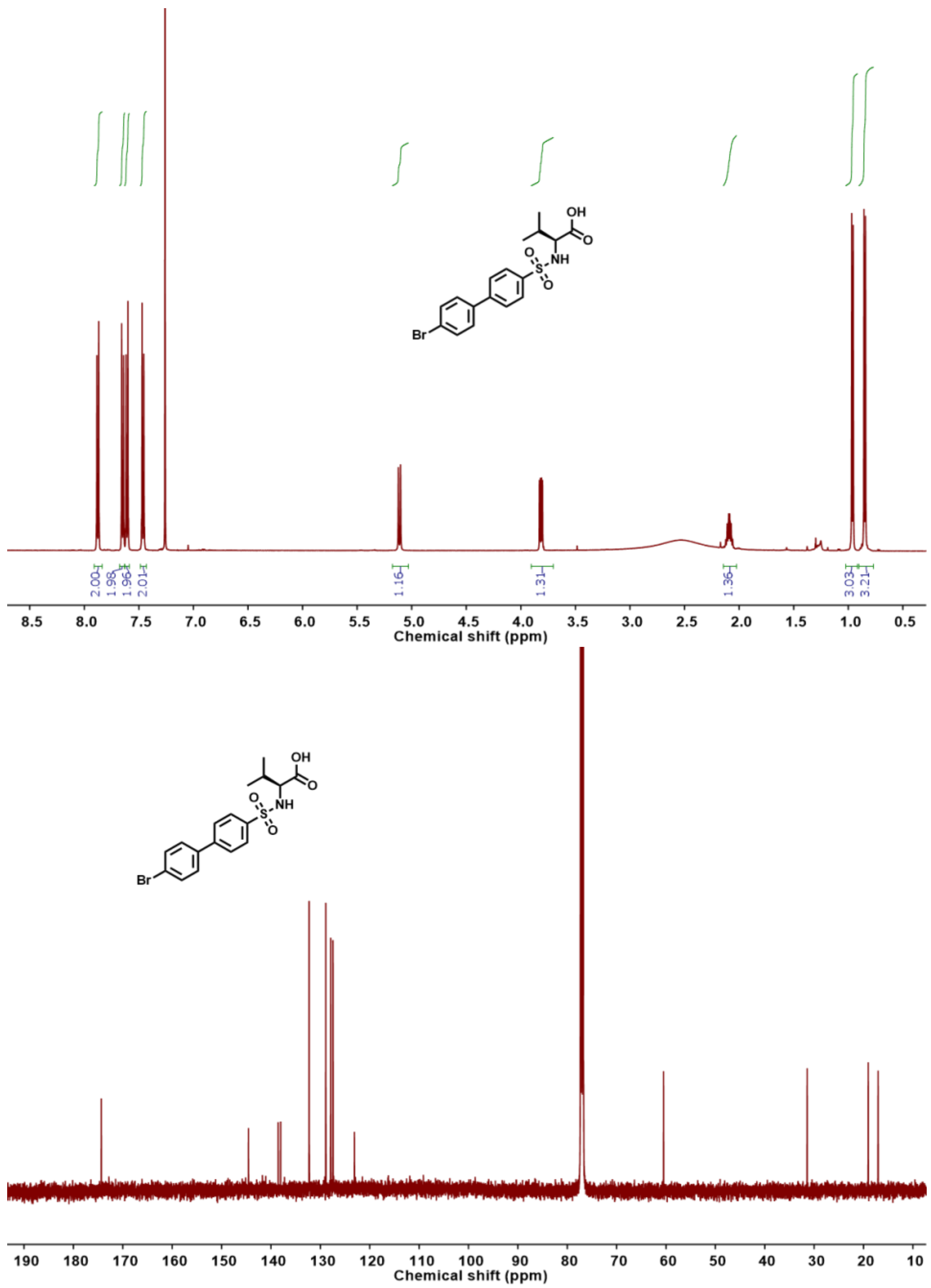


**Figure S11.** $^{1}H$ (top) and $^{13}C$ (bottom) NMR spectra of PD166793 in $CDCl_3$.

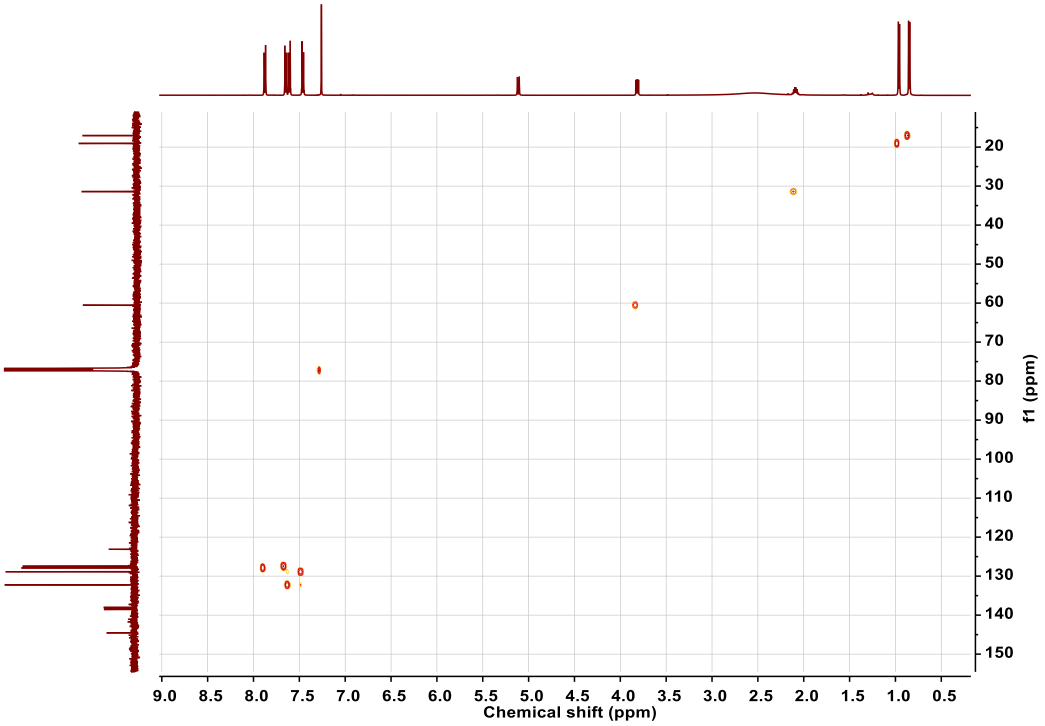


**Figure S12.** HSQC spectrum of PD166793 in $CDCl_3$.

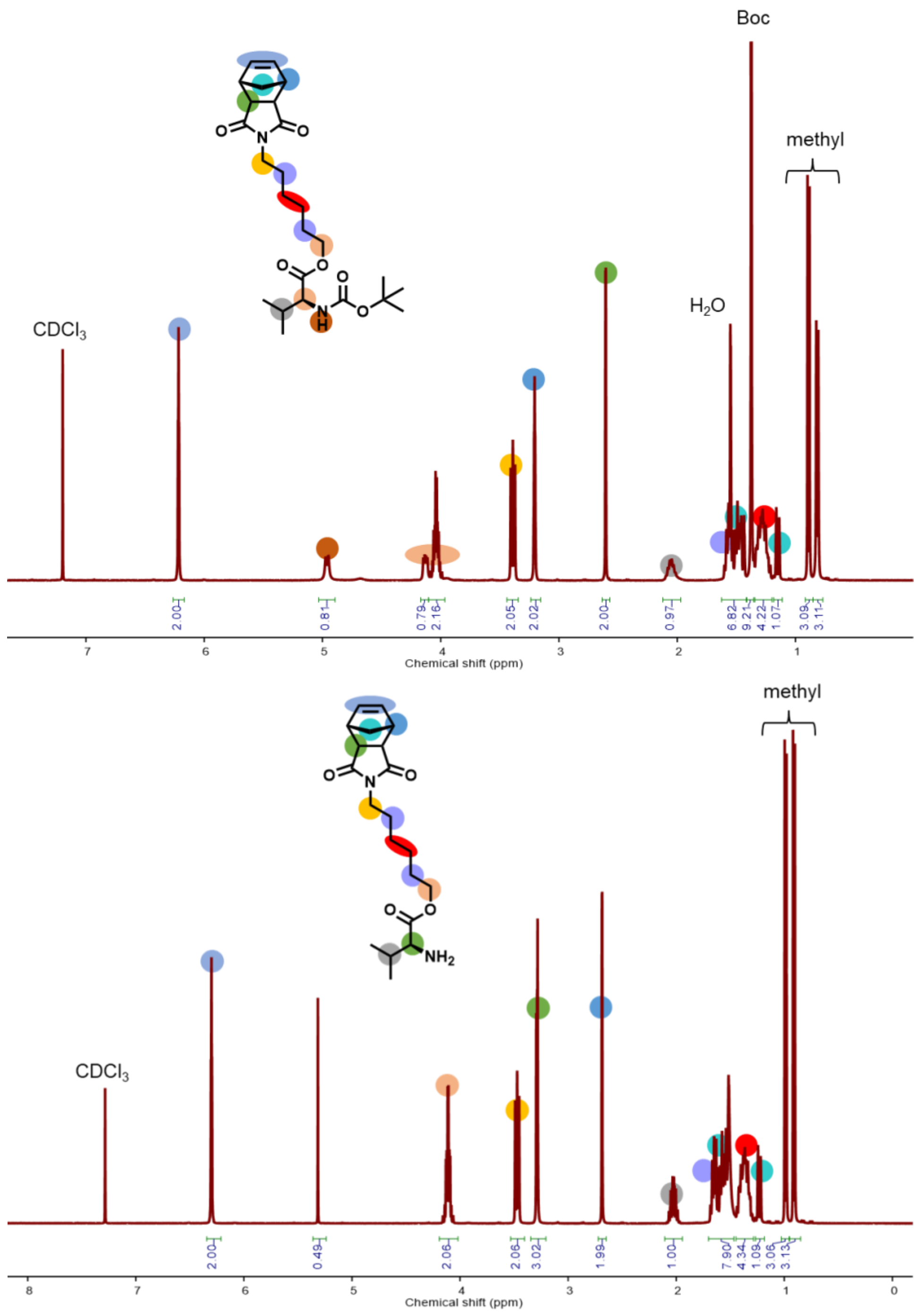


**Figure S13.** $^1$H NMR spectra of NorHex-Boc-Valine (top) and NorHex-Valine (bottom) in CDCl3.

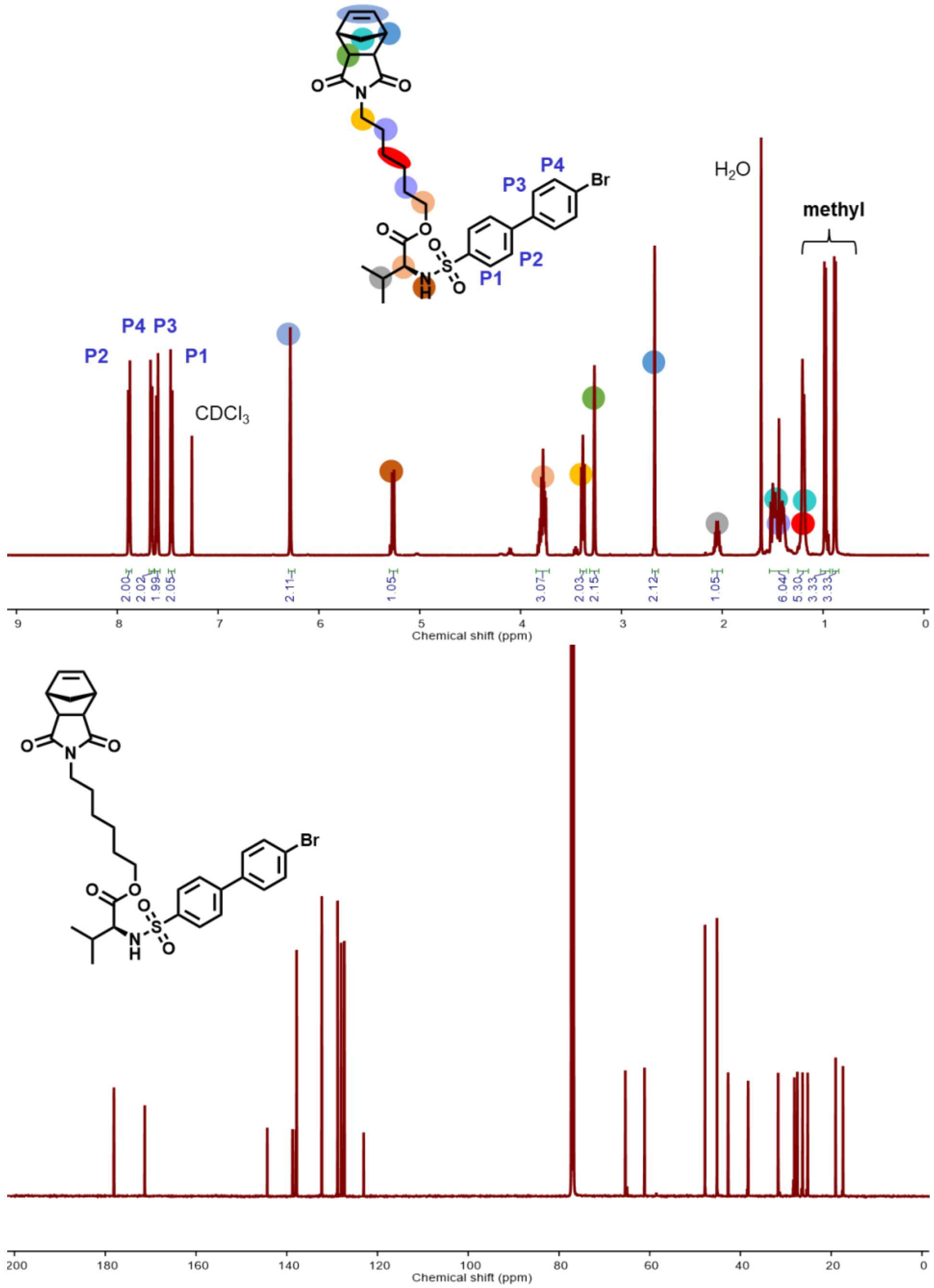


**Figure S14.** $^{1}H$ (top) and $^{13}C$ (bottom) NMR spectra of NorMMPi in $CDCl_3$.

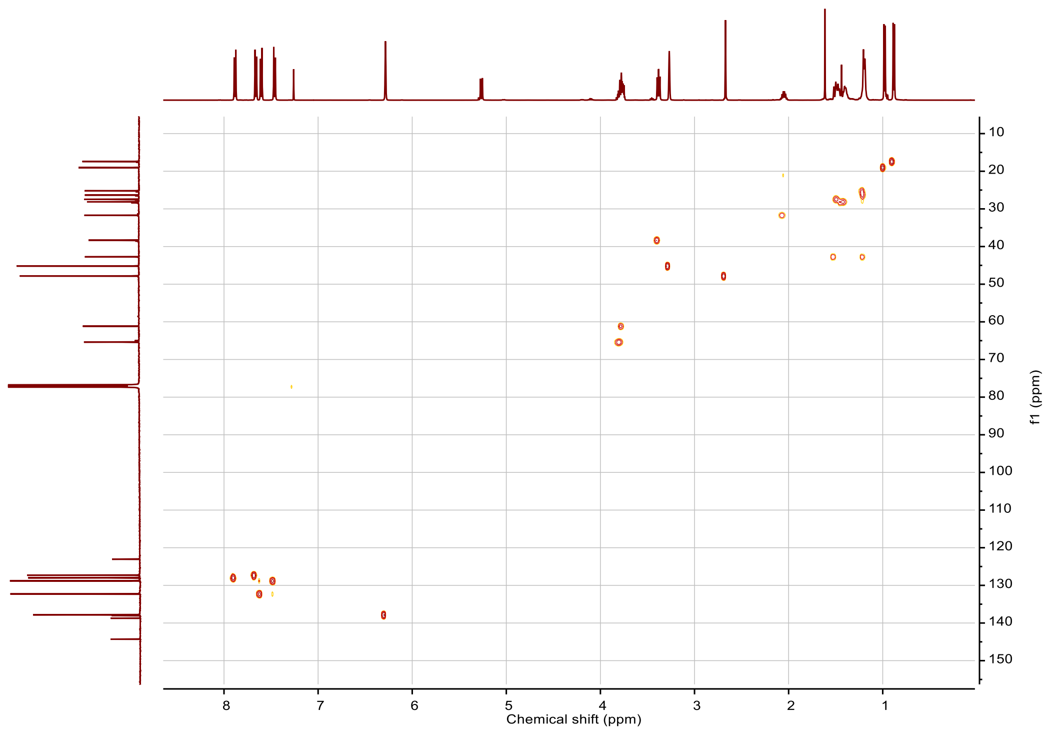


**Figure S15.** HSQC spectrum of NorMMPi in $CDCl_3$.
